# Phenology of 50 tree species across 9 years in a South Asian tropical rainforest indicates complex influence of climate, traits, and phylogeny

**DOI:** 10.64898/2026.02.14.705701

**Authors:** A. P. Madhavan, Srinivasan Kasinathan, Akhil Murali, K. B. Sonia, G. Moorthi, T. Sundarraj, R. Rajesh, Divya Mudappa, T. R. Shankar Raman

**Affiliations:** Nature Conservation Foundation, 1311, 12th A Main, Vijayanagar 1st Stage, Mysore 570017, Karnataka India

**Keywords:** tropical rainforest, signal processing, phenological phase, climate correlates, phylogenetic signal

## Abstract

In relatively aseasonal tropical rainforests, few studies have explored long-term phenological patterns of a high diversity of tree species in relation to climate, phylogeny, and functional traits. In these systems, short-duration seasonal pulses of irradiance and water deficit are expected to provide narrower windows for leafing and flowering and wider windows for fruiting across prolonged wet seasons, potentially mediated by functional traits and phylogenetic relatedness. Here, we document leafing, flowering, and fruiting phenology of 50 tree species (920 – 1077 trees, 10 – 42 trees/species) monitored monthly over an 9-y period (2017 – 26) in a relatively aseasonal south Asian tropical rainforest in the Anamalai Hills, Western Ghats, India. We examined correlations between climatic variables (irradiance, daylength, temperature, precipitation) and tree phenology and Mantel correlations among similarity in monthly phenology, functional traits (wood density, seed size, and maximum height), and phylogenetic relatedness. We then investigated the phylogenetic signal of phenological traits (frequency, amplitude, duration, and peak month) using Pagel’s λ. Leaf flushing and flowering showed distinct seasonality and negative associations with daylength and precipitation, whereas fruiting showed greater temporal spread and weaker associations with climate. Functional traits or phylogeny did not significantly influence leaf flushing and flowering, whereas dissimilarity in fruiting was correlated with phylogenetic distance and peak fruiting month showed a significant phylogenetic signal (λ = 0.96). The results indicate that in relatively aseasonal tropical rainforests, proximate climatic cues more strongly influence leaf flushing and flowering, whereas phylogenetic constraints affect timing of fruiting and may cause lineage-specific vulnerabilities to climate change.

## Introduction

Phenological systems of tropical trees are markedly complex, exhibiting greater spatial and temporal variability than the more predictable and seasonal cycles of temperate ecosystems (Wright 1996; Lex Engel and Martins 2005; Davis et al. 2022). Much of this complexity stems from the high climatic variability and weak annual seasonality of many tropical regions, especially rainforests near the equator, where resource limitation is minimal and productivity is sustained year-round (Wright 1996; Corlett and Lafrankie 1998; Zimmerman et al. 2007). The combination of unpredictable weather, spatial heterogeneity, and species-specific biotic factors creates low generalisability and predictability of phenological timing in these aseasonal forests (Corlett and Lafrankie 1998; Lex Engel and Martins 2005; Cleland et al. 2007; Morellato et al. 2013; Fitchett et al. 2015; Ford et al. 2016; Dunham et al. 2018; Davis et al. 2022). Even within a single landscape, co-occurring species can exhibit divergent phenological rhythms, including circannual, sub-annual, irregular, or supra-annual cycles, with low temporal synchrony (Lex Engel and Martins 2005; Brearley et al. 2007; Helm et al. 2013; Dunham et al. 2018; Cortés-Flores et al. 2019). Explaining this variability is essential for understanding how relatively aseasonal tropical systems function and will respond to climate change.

Above all, abiotic climatic cues impose the strongest control on phenological expression providing the environmental windows within which species-specific strategies and biotic interactions unfold (van Schaik et al. 1993; Wright 1996; Günter et al. 2008; Seghieri et al. 2012). In relatively aseasonal tropical rainforests, episodes of heightened irradiance act as the principal limiting factor, given that dense cloud cover occurs during more than nine months of the annual cycle (van Schaik et al. 1993; Graham et al. 2003; Hamann 2004). Consequently, phenophases such as leaf flushing and flowering, which depend on high irradiance to sustain their increased photosynthetic requirements, tend to concentrate within these short favourable periods, thereby promoting pronounced intra and interspecific synchrony and clustering (van Schaik et al. 1993; Wright and van Schaik 1994; Hamann 2004; Zimmerman et al. 2007; Calle et al. 2010; Manoli et al. 2018). In contrast, fruiting is shaped primarily by moisture availability, photoperiod, and temperature (Günter et al. 2008; Seghieri et al. 2012; Verma et al. 2022), which are factors that remain relatively high or uniform across much of the year in tropical rainforests (van Schaik et al. 1993; Wright 1996; Zimmerman et al. 2007). This creates a broad fruiting window allowing species to express multiple phenological strategies that may be temporally diffuse (Wright and Calderon 1995; Brearley et al. 2007; Visser et al. 2010; Morellato et al. 2013; Dunham et al. 2018; Cortés-Flores et al. 2019). Even staggered fruiting can follow clear organization: in Central American dry forests, anemochoric species peak in early dry season, endozoochorous species from pre-wet to early wet season, and autochorous and epizoochorous species later in wet season (Cortés-Flores et al. 2019). These multiple fruiting optima are underpinned by physiological differences in fruit development time, seed size and plant growth form (Buckley and Kingsolver 2012; Helm et al. 2013; Cortés-Flores et al. 2019).

Ultimately, the diversity of phenological strategies may reflect evolutionary adaptations conserved within phylogenies, either through plant physiological or functional traits or the direct conservation of phenological timing, frequency, or duration (Wright and Calderon 1995; Willis et al. 2008; Buckley and Kingsolver 2012; Davies et al. 2013). Evidence for phylogenetic signals are clear for some functional traits such as seed size or development time, which can then mediate phenological strategy (Cortés-Flores et al. 2019). However, phenological patterns of species can be directly conserved in phylogenetic lineages; for example, globally, phylogenetically related species tend to initiate flowering and fruiting at more similar times, even when they occur in different biomes or on different continents (Davies et al. 2013). In the context of tropical rainforest tree phenology, it remains unclear whether multiple phenological strategies within the same landscape are driven directly by phylogenetic constraints, indirectly through functional traits, or by functional traits that operate independently of phylogeny due to convergent evolution (Buckley and Kingsolver 2012; Davies et al. 2013; Helm et al. 2013). Resolving the relative contributions of phylogeny, functional traits, and the degree of internal vs external timekeeping mechanisms on tropical rainforest phenology can help predict how species will respond to shifting climatic cues (Willis et al. 2008; Davis et al. 2010; Buckley and Kingsolver 2012; Helm et al. 2013) and map the potential impacts of climate change on tropical forest dynamics and ecosystem functioning.

In the present study, we investigate how climate, plant physiological traits, and phylogenetic relatedness shape phenological patterns of leafing, flowering, and fruiting in 50 rainforest species monitored over 9 years within tropical rainforests of the Western Ghats mountain range, India. Specifically, we asked two main questions: (1) How do key climatic variables such as irradiance, temperature, humidity, precipitation and photoperiod correlate with vegetative and reproductive phenological patterns of tropical tree species in a relatively aseasonal south Asian tropical rainforest? (2) Are interspecific differences in vegetative and reproductive phenology better explained by functional trait variation or by phylogenetic relatedness, and does this influence differ across components of phenological expression including frequency, timing, intensity and duration? For Question 1, we hypothesized that leaf flushing and flowering are regulated primarily by irradiance-driven increases in photosynthetic carbon gain, whereas fruiting phenology is governed by moisture availability and photoperiod cues associated with seed maturation and germination. We predicted that leaf flushing and flowering would mainly peak during the limited high-irradiance months resulting in higher synchrony and shorter durations of these phenophases, while fruiting will be more staggered across the year due to its longer developmental period and weaker dependence on temporally limited climatic conditions. For Question 2, we evaluate whether interspecific phenological similarity is unrelated to either functional traits or phylogenetic relatedness or, alternatively, is driven independently by either functional traits or phylogenetic relatedness, or by both factors.

## Methods

### Study area

The study was conducted within the Anamalai hills, a significant conservation landscape that is part of the Western Ghats global biodiversity hotspot in India (Kumar et al. 2004). The main study area (Figure 1) was the Anamalai Tiger Reserve (ATR, core area: 958 km^2^, 10.2203° – 10.5588° N, 76.8163° – 77.4187° E) and the adjoining Valparai Plateau (220 km^2^, 10.2424° – 10.3877° N, 76.8287° – 77.0523° E). Although ATR contains a diversity of natural vegetation types, the study focused on mid-elevation (700 – 1400 m asl) tropical wet evergreen rainforests of the *Cullenia exarillata – Mesua ferrea – Palaquium ellipticum* floristic type (Pascal 1988; Pascal et al. 2004) that formed the main natural vegetation type on the Valparai Plateau and adjoining parts of ATR.

**Fig. 1.**
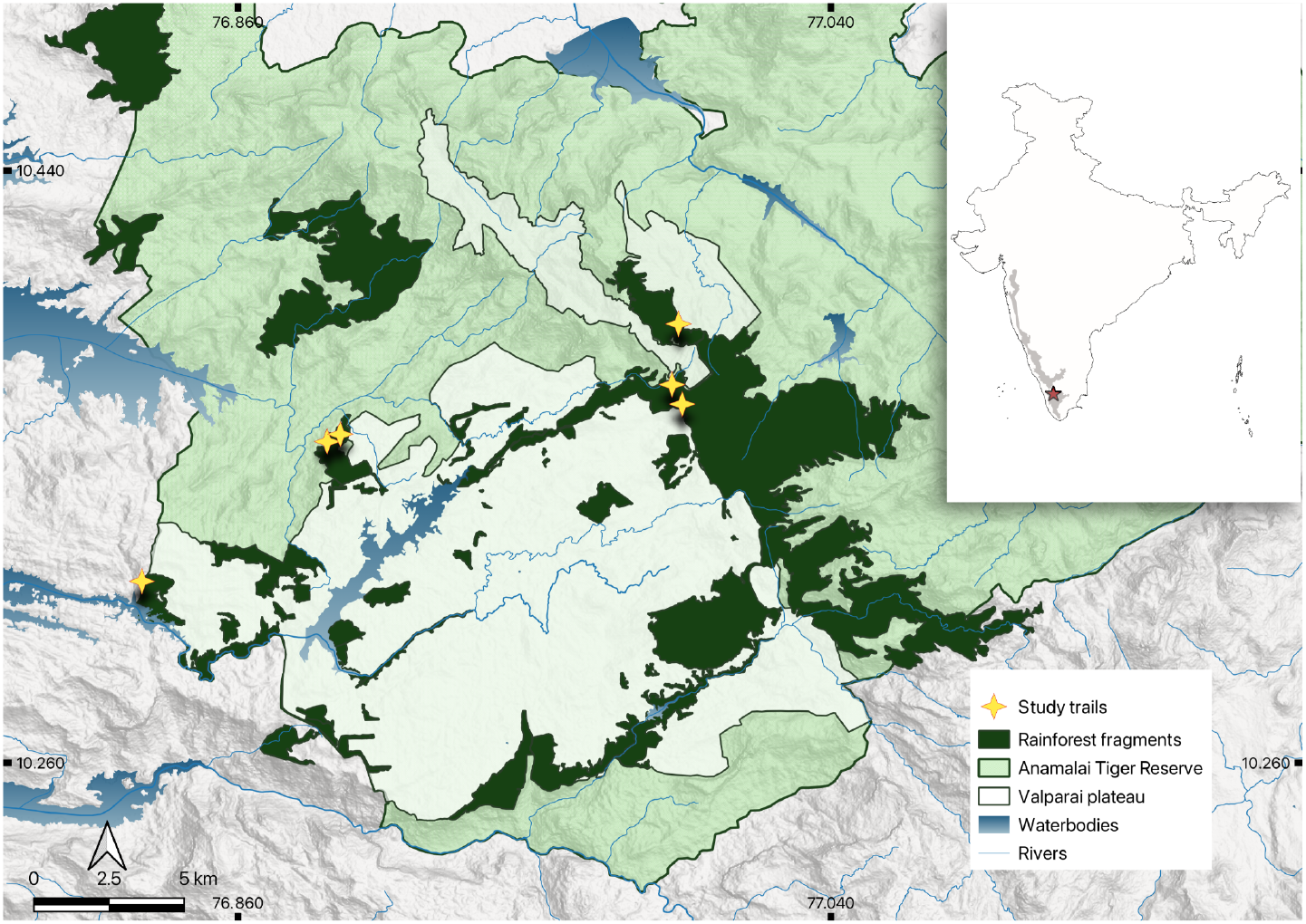
Map of study area including the study trails, terrain, rainforests fragments, protected area boundaries and waterways.

Climate data (rainfall, temperature, radiance, photoperiod and relative humidity) for the study area were recorded using a local electronic weather station (HOBO U30 Station, Onset Computer Corporation) maintained by the Anamalai Rainforest Research Station at Iyerpadi (10.363725° N, 76.977628° E, 1234 m asl) on the Valparai Plateau. Weather station data between 2014 – 2025 visualised in Figure 2 summarises the seasonal variation in climate patterns. The study area receives a mean annual rainfall of approximately 2650 mm, mainly between June and September (southwest monsoon). Additional rainfall occurs between October and December (northeast monsoon) and as pre-monsoon thunderstorms in April and May, resulting in only a brief dry period from January to March. Irradiance ranges from 350 μmol m^−2^ s^−1^ between February to March to a low of 180 μmol m^−2^ s^−1^ during the peak wet season from July to September. Average relative humidity is high (85.8%), peaking at 96–98% from June to August and dropping to 69–75% from January to March. Temperature is distinctly bimodal (Figure 2b) with a higher peak during April – May and a second lower peak after the SW monsoon in October – November. Mean annual temperature is 20.8°C, with the warmest conditions in April (23.5°C) and cooler periods (<19.7°C) in December and July – August. Photoperiod averages 12 hours, varying modestly from a minimum of 11.61 hours in December to a maximum of 12.5 hours in April–May. Overall, the site exhibits limited seasonality in temperature, humidity, and photoperiod, with most climatic seasonality driven by irradiance and rainfall patterns.

**Fig. 2.**
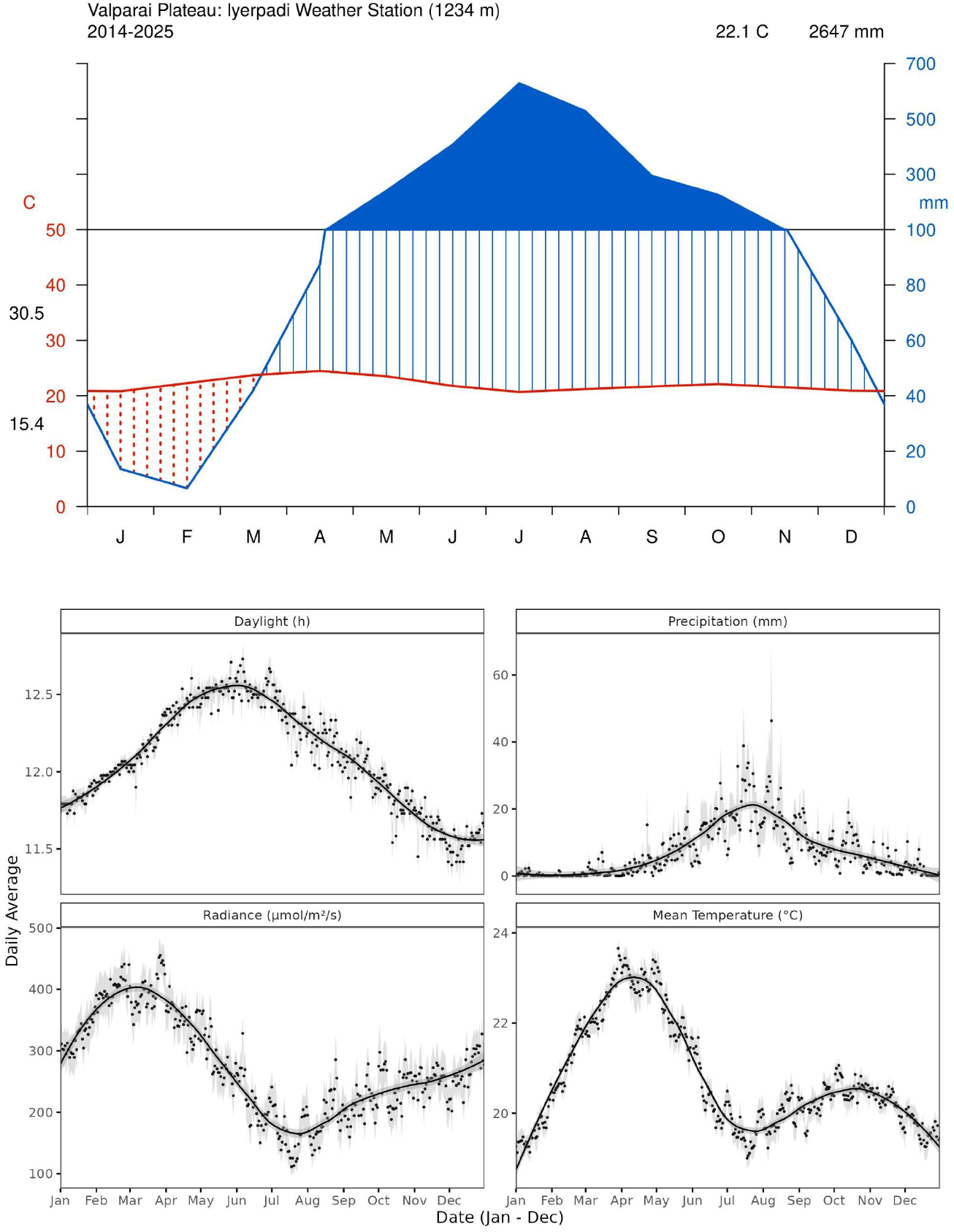
Climate summary for the study area (2014 – 2025): (a) Walter-Leith Diagram (b) annual pattern of key weather parameters (lines shows loess fit of daily means at span = 0.4; points are daily means; grey shaded area shows ±1 SE of daily mean.)

### Study species

We focused on 50 tree species of (920 – 1077 individuals overall), with 10 – 42 trees per species across 25 families monitored monthly over 9 years (108 months) from March 2017 – February 2026. These 50 species made up about 70% of the individuals in the tree community along the surveyed transects. Dominant families within our study sample included Lauraceae (7 species); Euphorbiaceae (6 species) followed by Meliaceae and Phyllanthaceae (4 species each).

### Field methods

Seven phenology trails (total length 2.9 km; 265 m – 575 m length) were established and sampled monthly from March 2017 along existing access routes in ATR (6 trails) and the Valparai Plateau (1 trail), spanning the elevational range of 700 m – 1405 m and floristic range of the mid-elevation tropical wet evergreen rainforest type. One trail at 700 m elevation in ATR was discontinued from October 2020 due to reasons of logistics, leaving the remaining six spanning 790 m – 1405 m. Within each trail, all trees ≥30 cm girth at breast height (gbh, 1.3 m) located within 5 m on either side of the edge of each trail were identified to species, measured for girth (using tape measure) and tree height (using a rangefinder), and mapped with geographical coordinates obtained using a Garmin eTrex GPS handset. For the present study, we used data from all tree species that were represented by at least 10 individuals, capping the number of monitored individuals of more common species at 40 individuals. To complement a parallel study on threatened trees, we added 97 individuals of six threatened tree species from along or near the phenology trails in March 2021 (to achieve a minimum of 20 individuals per species). Trees were monitored for phenology at the beginning of each calendar month: from March 2017 onwards for the 920 trees of 44 species, and from March 2021 onwards for the 97 individuals of the six threatened species.

Five phenophases—leaf flush, flower buds, flower open, unripe fruits, and ripe fruits—were observed and recorded for each individual tree by two observers using 8×42 binoculars. Through a complete visual scan of the crown, phenophase intensity was scored 0 – 4 based on the percentage of canopy estimated to have the phenophase (0: 0% canopy; 1: 1 – 25%; 2: 26 – 50%; 3: 51 – 75%; 4: 76 – 100%). Dead individuals were not replaced, so sample sizes for some species decreased slightly over time. Damage or death was recorded, and individuals were monitored for resprouted shoots for 12 – 24 months.

### Data analysis

For the purposes of this study, we used data only from the 6 transects surveyed across all 9 years. From the phenological data we selected 3 phenophases of interest: leaf flush, flower open, and ripe fruit. For each species, the proportion of individuals expressing a particular phenophase in a given month was calculated and used as the fundamental phenological parameter of interest. Based on the pattern of phenophase expression over time, we analysed phenological patterns using a signal processing approach by comparing neighbour value deltas using functions of R package pracma (Borchers, 2011). The start of a peak was identified as the last month before the phenophase expression (proportion) increased, with the peak then defined as the last month before the value decreased for two consecutive months or before it levelled off. To identify peaks along the time series, we imposed a minimum period between peaks as 4 months to identify subsequent independent peaks, taking this as a reasonable assumption given reproductive biology of rainforest trees and seasonal variation. Using this approach, we calculated the duration, frequency, amplitude, and peak month for each species over time based on the proportion of individuals expressing a given phenophase, repeated for all 3 phenophases. Duration of phenophase (months/year) was calculated as 12 × proportion of months phenophase expression exceeded the absolute minimum recorded across the nine years for 44 species and 5 years for the 6 threatened species. Frequency was defined as the number of distinct peaks, and peak month corresponded to the months during which those peaks occurred. Amplitude was the proportion value associated with the identified peak month. Based on the calculated frequency, species were classified into five categories: supra-annual (frequency ≤0.7 peaks year^−1^), annual (0.7 – 1.3 peaks year^−1^), sub-annual (1.3 – 1.8 peaks year^−1^), and semi-continuous or irregular (>1.8 peaks year^−1^) (modified from Newstrom et al. 1994).

Using the weather station data, monthly rainfall was calculated as a monthly total rainfall received, while daily temperature, humidity, and irradiance were averaged by month, and photoperiod was calculated by taking the number of daylight hours indicated by photosynthetically available radiation (PAR > 1.2 μmol m^−2^ s^−1^) each day and averaging by month. Prior to averaging, gaps in daily data, due to temporary sensor failure or equipment malfunction, were filled with a 7-day rolling mean of the 2014 – 25 values centred on the corresponding date.

We assessed the role of climate in driving phenological patterns within this relatively aseasonal rainforest system using monthly time-series cross-correlation and cross-covariance analyses between climatic variables and the phenophase expression for each species. These analyses were performed at a one-month lag to evaluate whether changes in irradiance, temperature, precipitation, or photoperiod in the preceding month were associated with shifts in proportion of individuals expressing a phenophase in the beginning of that month. For each species, positive cross-correlation values were interpreted as positive cues for phenophase expression (e.g., higher irradiance leading to greater leaf flushing intensity), whereas negative values indicated inhibitory cues (e.g., declining humidity preceding increases in flowering). The magnitude of the correlation reflected the strength of the association, with values closer to ±1 indicating stronger climate – phenophase relationships. This approach allowed us to identify which climatic variables most strongly shaped the timing and patterns of expression of each phenophase across species.

To assess whether interspecific differences in monthly phenology were better explained by differences in functional traits or by phylogenetic relatedness, we compared species-level dissimilarity matrices in phenology, functional traits, and phylogeny. A pairwise phenological dissimilarity matrix among species based on their monthly phenophase values across the 108-month time series was constructed using Euclidean distance using the ‘vegdist()’ function in R package vegan (Oksanen et al. 2025). Using this method, we computed separate dissimilarity matrices for leafing, flowering, and ripe fruits. Functional traits, including wood density, seed size, and maximum height, were obtained from the Zenodo database (Osuri et al. 2022). These traits were used to construct a pairwise functional dissimilarity matrix among species using Euclidean distance. Seed size categorised as small (<1 cm), medium (1 – 3 cm), and large (>3 cm) were taken as 1, 2, and 4 for calculating the distance matrix. Phylogenetic dissimilarity was quantified using pairwise branch lengths derived from a resolved species phylogeny generated with the V.PhyloMaker2 R package using the phylo.maker() function which grafts species from the GBOTB (global taxonomic backbone tree of vascular plants) megatree (Jin and Qian 2023). We used Mantel tests to assess relationships between dissimilarity matrices taking significant (*p* < 0.05) positive Mantel correlation coefficients (*r*) to indicate that species that differ more strongly in their phenology also differ more strongly in their traits or phylogenetic distances. Significance of the Mantel *r* coefficient was assessed using 10,000 permutations.

Lastly, we quantified the extent to which phenological traits exhibit phylogenetic signals by estimating Pagel’s λ for each phenological trait (frequency, duration, peak month, amplitude) across all four phenophases using the ‘phylosig()’ function in phytools R package (Revell 2012). Pagel’s λ quantifies the extent to which closely related species resemble each other in a given trait. Values of λ close to 1 indicate strong phylogenetic dependence consistent with Brownian-motion evolution, whereas values of λ near 0 indicate that phenological traits show little phylogenetic structuring and are evolutionarily labile (Pearse et al. 2025). Significance of the estimated λ was assessed using a likelihood ratio test that compares the log-likelihood of the fitted model to that of a model with λ constrained to zero (no phylogenetic signal). The resulting test statistic is evaluated against a χ^2^ distribution with one degree of freedom, providing a *p*-value indicating whether the observed phylogenetic signal differs from random expectations.

## Results

### Phenophase expression

Species frequency of phenological peaks exhibited a broad range (0.1 – 2 peaks per year), yet median and mean values clustered near 1.0 for all phenophases (Figure 3). However, frequency distributions differed significantly among phenophases, with leaf flushing skewed toward biannual patterns (1.5 – 2 peaks year^−1^) and fruit ripening skewed toward supra-annual patterns (<0.6 peaks year^−1^). Phenological amplitude varied substantially, averaging highest in leaf flushing (0.87), intermediate in flowering (0.5), and lowest in ripe fruiting (0.35). Phenophase duration also varied, with leaf flushing persisting longest (mean: 10.8 months), followed by flowering (mean: 5.5 months) and ripe fruiting (mean: 4.8 months). Ripe fruiting exhibited large variation (interquartile range: 2.4–7 months), with numerous outliers present in both flowering and fruiting. Months of peak phenophase expression as indicated by interquartile range of values across the 50 species was February – May for leaf flushing and March – June for flowering, both skewed toward earlier months. In contrast, fruit ripening peaked later (interquartile range: May – September) and exhibited greater variation with broader tails on both sides of the distribution, indicating greater interspecific variation in timing.

**Fig. 3.**
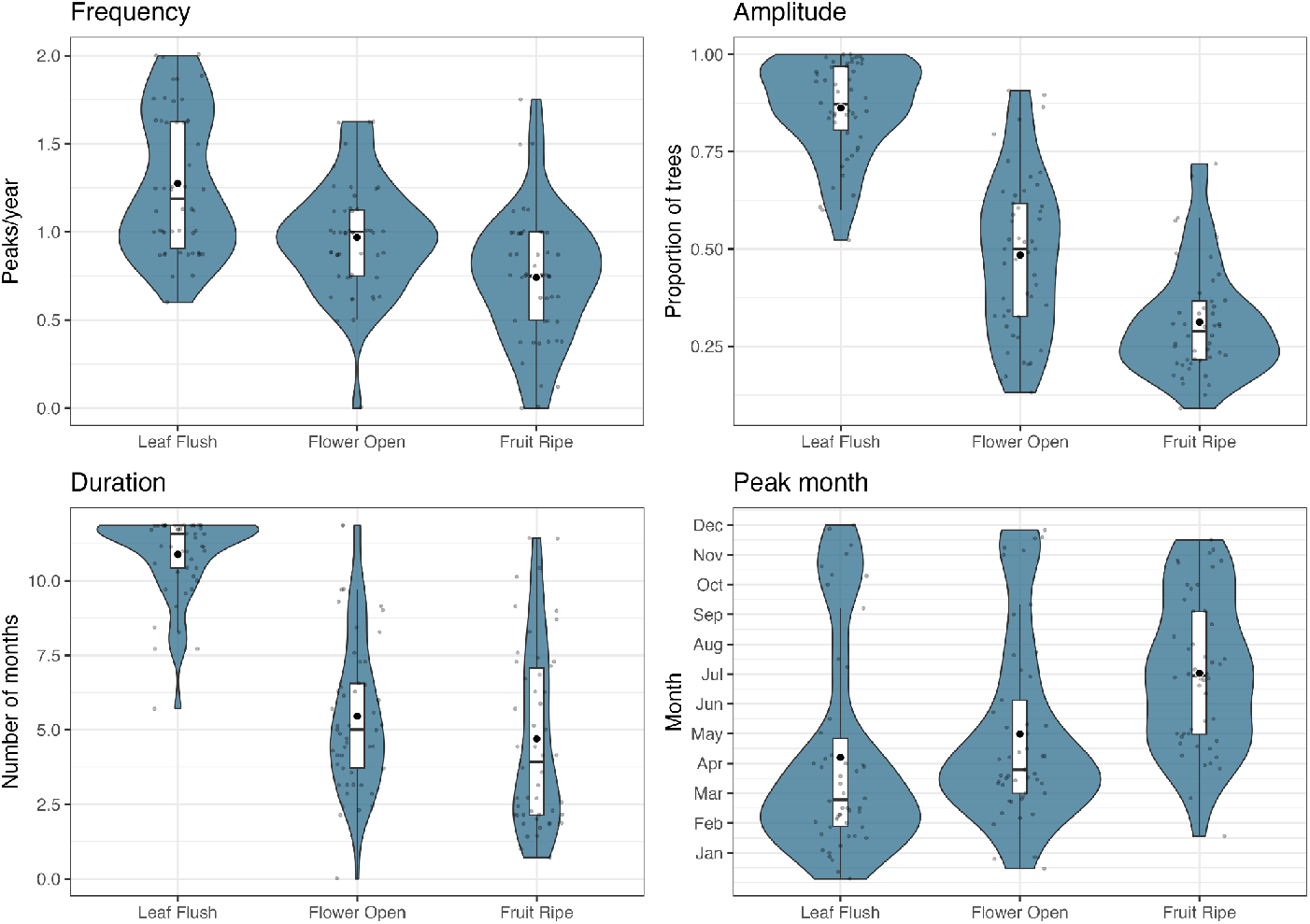
Phenological trait variation across phenophases. Box plots represent the median and interquartile range, violin plots show the kernel density distribution of values, black dots indicate means, and faded blue dots represent individual species values.

### Climate – phenophase cross-correlations

Climate – phenology relationships differed strongly among phenophases (Figure 4). Leaf flushing and flowering exhibited clear negative associations with precipitation and daylength in the previous month, with 30 species (out of 50) showing significant negative cross-correlations for leaf flushing, and 35 and 30 species for flowering (|*r*| > 0.19, df = 105, P < 0.05), respectively. Additionally, in leaf flushing 21 species showed significant positive association with irradiance in the previous month. In ripe fruiting, while 21 species showed significant positive associations with temperature and 19 with daylength, a larger number of species had non-significant correlations across all climate variables (Figure 4).

**Fig. 4.**
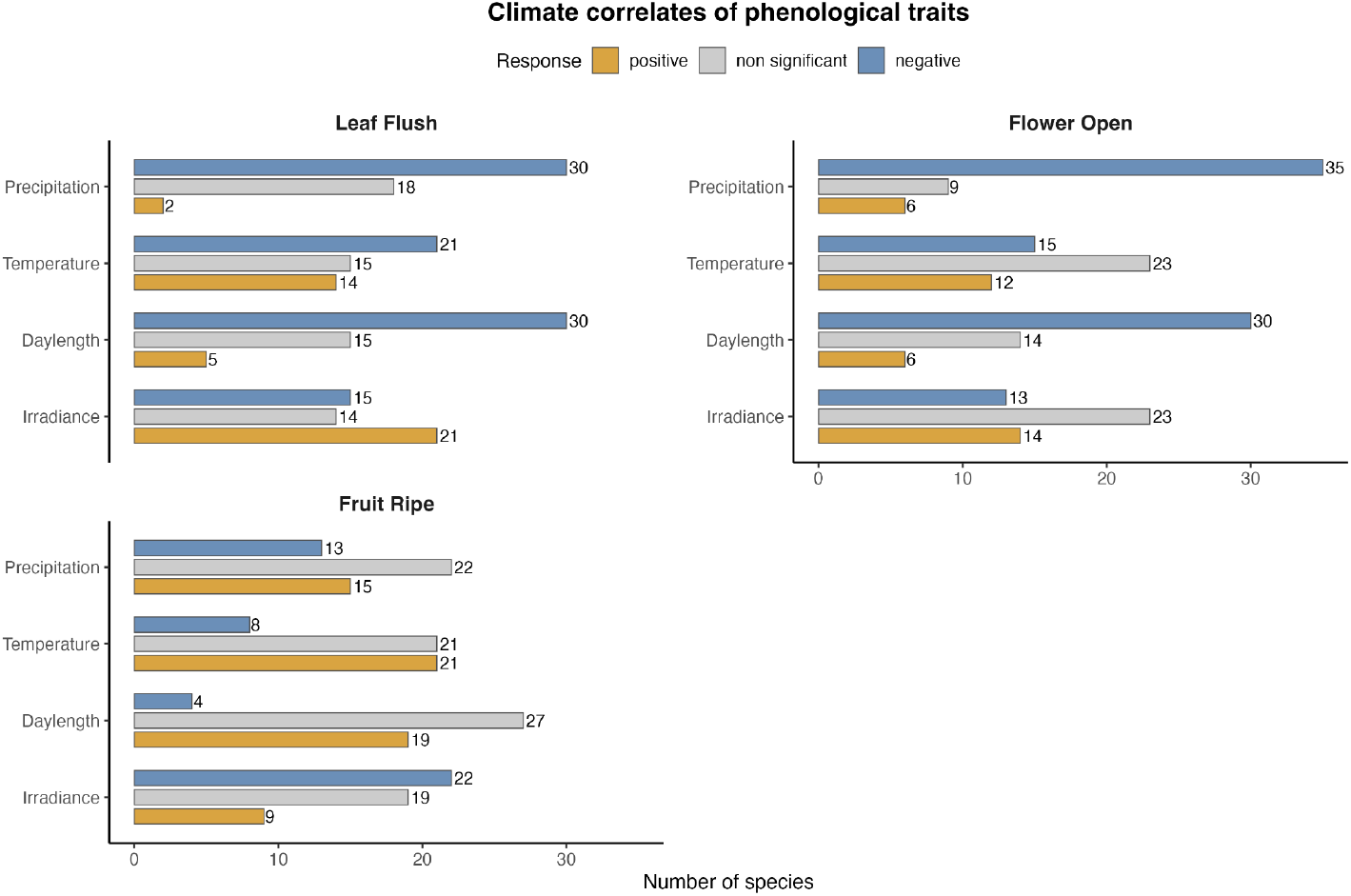
Cross-correlations between monthly phenophase expression and four key climatic variables measured at lag 1 (previous month); bars indicate the number of species showing significant (Pearson’s r, P < 0.05) positive or negative relationships, and the number of species where the effects were non-significant (P > 0.05).

### Influence of phylogeny and traits

Mantel tests revealed little evidence for phylogenetic or trait-based structuring of phenology across most phenophases (Table 1). Leaf flushing and flowering showed no significant associations between phenological similarity and either phylogenetic distance or functional trait dissimilarity (Mantel *r* = −0.08 to 0.04, *P* > 0.17) (Table 1). Fruiting phenology showed a weak but significant phylogenetic signal (Mantel *r* = 0.13, *P* = 0.01), while trait – phenology relationships remained non-significant (Table 1). Dissimilarity in functional traits was also correlated to phylogenetic distance (Mantel *r* ≃ 0.00, *P* = 0.50).

**Table 1.**
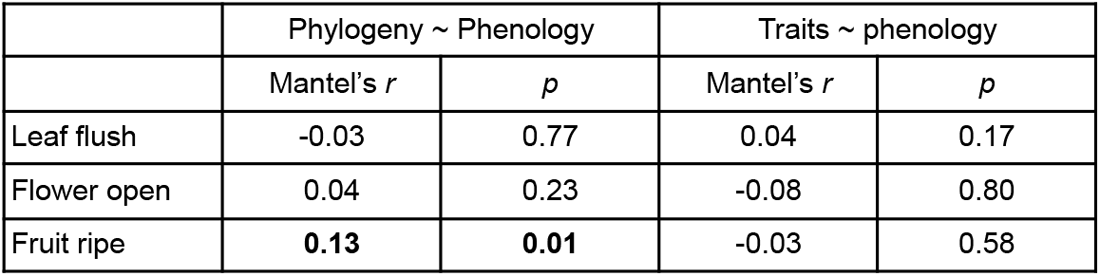
Results of Mantel tests for relationships between phylogenetic distance, functional trait distance, and phenological distance across three different phenophases in the 50 tree species.

|  | Phylogeny ~ Phenology |  | Traits ~ phenology |  |
| --- | --- | --- | --- | --- |
| | Mantel's $r$ | $p$ | Mantel's $r$ | $p$ |
| Leaf flush | -0.03 | 0.77 | 0.04 | 0.17 |
| Flower open | 0.04 | 0.23 | -0.08 | 0.80 |
| Fruit ripe | **0.13** | **0.01** | -0.03 | 0.58 |

Overall most phenological traits had no phylogenetic signal (Table 2). Among phenological traits, the peak month of fruit ripening exhibited a strong and significant phylogenetic signal (λ = 0.96, *P* = 0.03), followed by fruiting duration (λ = 0.75, *P* = 0.04) (Table 2).

**Table 2.**
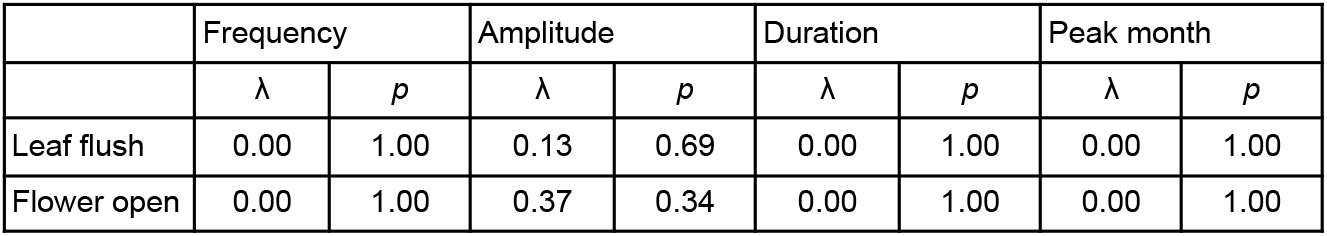
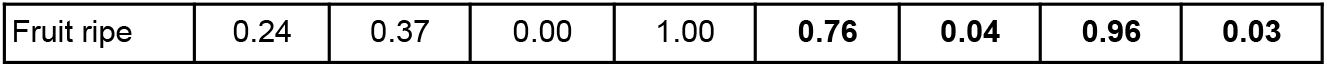
Pagel’s λ for the four phenological traits across three phenophases for the 50 tree species (statistical significance of the *p* value based on likelihood ratio tests).

|  | Frequency |  | Amplitude |  | Duration |  | Peak month |  |
| --- | --- | --- | --- | --- | --- | --- | --- | --- |
| | $\lambda$ | $p$ | $\lambda$ | $p$ | $\lambda$ | $p$ | $\lambda$ | $p$ |
| Leaf flush | 0.00 | 1.00 | 0.13 | 0.69 | 0.00 | 1.00 | 0.00 | 1.00 |
| Flower open | 0.00 | 1.00 | 0.37 | 0.34 | 0.00 | 1.00 | 0.00 | 1.00 |
| Fruit ripe | 0.24 | 0.37 | 0.00 | 1.00 | **0.76** | **0.04** | **0.96** | **0.03** |

Species in families Lauraceae, Sapotaceae, and Myrtaceae tended to peak in pre-monsoon (April – May), Meliaceae and Phyllanthaceae in the first wet season (June – August) and Rutaceae, Euphorbiaceae, Ebenaceae, and Anacardiaceae in the second wet season (October – December) (Figure 5). Phylogenetic signal in ripe fruiting duration was also evident (Online Resource 1) with families like Meliaceae and Lauraceae tending to have shorter (<5 months) and others like Phyllanthaceae and Euphorbiaceae longer (>6 months) durations.

**Fig. 5.**
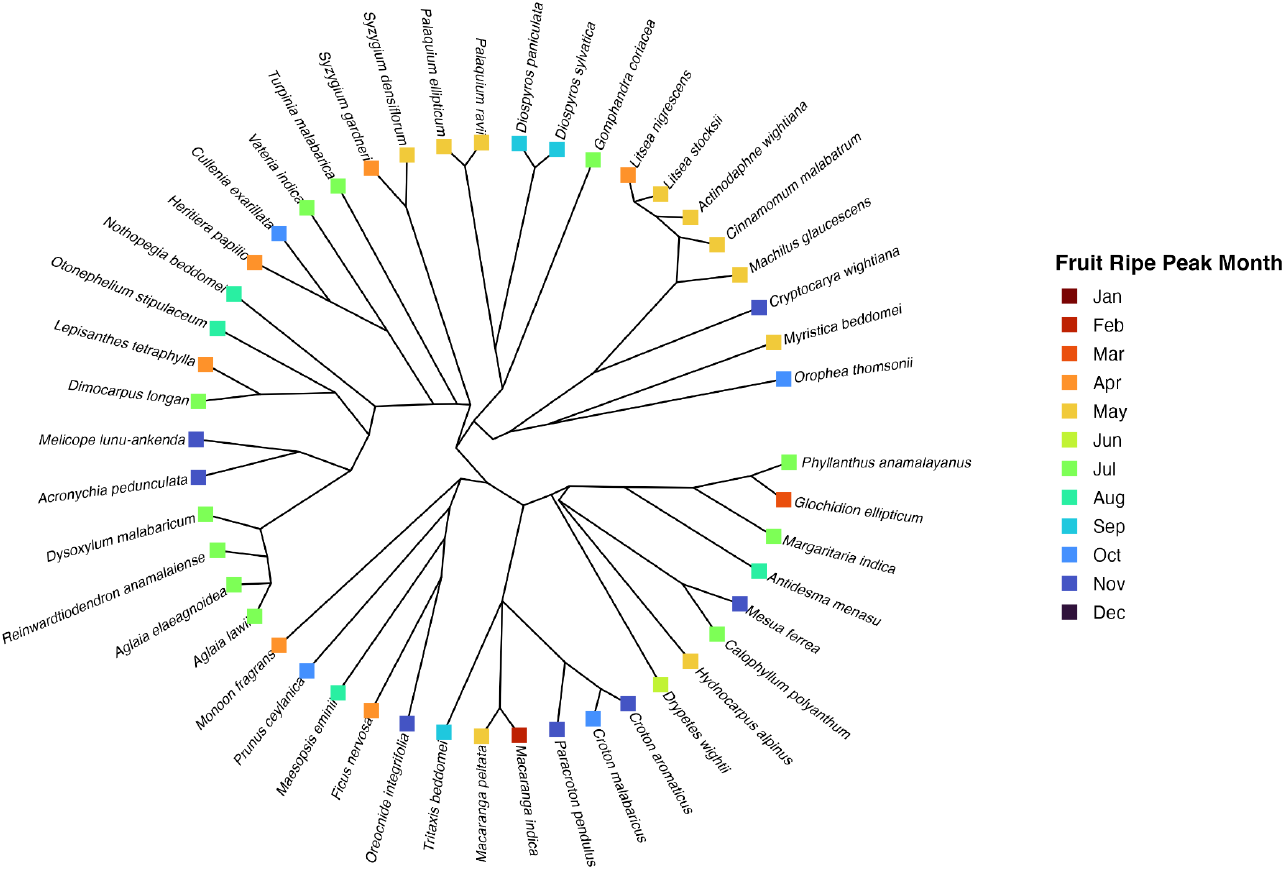
Phylogenetic signal in peak fruiting month. The phylogenetic tree shows evolutionary relationships among species, with branch lengths representing time since divergence. Colors indicate the peak month of ripe fruiting for each species.

## Discussion

We predicted that leaf flushing and flowering would peak during periods of high irradiance, resulting in high synchrony and short, well-defined phenological windows, whereas fruiting would be more temporally dispersed due to its longer and variable developmental period and weaker climatic influence. This prediction was more strongly supported for leaf flushing than for flowering. Leaf flushing peaked earlier in the year and had high amplitudes, which could indicate greater synchrony amongst individuals (Figure 3). Additionally, leaf flush showed consistent negative associations with precipitation and daylength, while also exhibiting positive associations with irradiance in 21 species (Figure 4). Flowering, although similarly seasonal and synchronous, showed weaker association with irradiance but had strong negative association with precipitation and daylength. This indicates that while canopy foliage renewal may have a stronger linkage to light availability cues (Wright and van Schaik 1994; Barone 1998; Zimmerman et al. 2007) in relatively aseasonal landscape, the absence of rainfall and associated cloud cover may be a stronger determinant of flowering windows (van Schaik et al. 1993; Graham et al. 2003; Singh and Kushwaha 2006; Günter et al. 2008; Eguchi et al. 2025). Dry periods with reduced cloud cover may provide optimal flowering conditions by minimizing flower damage from heavy rainfall (Lawson and Rands 2019) and by coinciding with warmer, sunnier days when insect pollinator activity is higher (Cruden 1972; Corbet et al. 1993; Peat and Goulson 2005). The stronger coupling of leaf flushing to irradiance may explain the higher incidence of biannual flushing, with favorable conditions for leaf production occurring during both pre-monsoon (March – April) and post-monsoon (November – December) periods in the southern Western Ghats (Padmakumari et al. 2022).

In contrast to leaf flushing and flowering, and in line with predictions, fruiting exhibited lower synchrony, broader peak timing distributions (Figure 3), and weaker climate – phenology relationships (Figure 4). Such patterns are consistent with studies showing greater temporal flexibility in fruiting, where weak abiotic constraints allow biotic factors to dominate phenological timing (van Schaik et al. 1993; Wright and Calderon 1995; Brearley et al. 2007; Morellato et al. 2013), particularly selection for optimal windows that maximise seed germination and seedling establishment. For many tropical tree species with short dormancy periods, fruiting is therefore timed to precede or coincide with the wettest part of the year, when early life-stage growth is most favourable (van Schaik et al. 1993; Grombone-Guaratini and Rodrigues 2002). However, this need not apply to all species, as interspecific differences in dispersal strategies, frugivore interactions, and associated resource competition trade-offs can shift the fruiting optima and duration for a species (van Schaik et al. 1993; Wright 1996; Wiens and Graham 2005). Dispersal mechanism, in particular, acts as an important temporal organizer: wind-dispersed species concentrate fruiting in the dry season to exploit optimal atmospheric conditions for long-distance dispersal (van Schaik et al. 1993; Griz and Machado 2001), mammal- and bird-dispersed species in the pre-monsoon and early monsoon to align with frugivore breeding seasons and peak activity, and gravity-dispersed species in the peak wet season when moisture facilitates fruit-splitting mechanisms (Cortés-Flores et al. 2019).

In Question 2, we examined the joint or independent influence of functional traits and phylogeny on phenological patterns using Mantel tests which indicated that interspecific phenological similarity was largely independent of both phylogenetic relatedness and functional trait similarity across leafing and flowering phenophases (Table 1). The absence of phylogenetic or trait-based structuring in leaf flushing and flowering suggests that these phenophases are evolutionarily labile and strongly shaped by environmental cues, allowing closely and distantly related species to converge on similar seasonal timing (CaraDonna and Inouye 2015; Zhang et al. 2017; Cortés-Flores et al. 2019). This decoupling from traits and phylogeny implies a higher degree of phenological plasticity, which may facilitate community-level synchrony driven primarily by local climate regimes for these variables (Wright and Calderon 1995; CaraDonna and Inouye 2015). In contrast, fruiting phenology showed a weak but significant phylogenetic signal, indicating that reproductive timing is more evolutionarily constrained, potentially reflecting lineage-specific strategies linked to seed development, dispersal, or establishment irrespective of similarities in traits and morphology (Davies et al. 2013). This interpretation is further supported by Pagel’s λ analyses, which revealed a strong phylogenetic signal in the peak month of fruit ripening and a moderate signal in fruiting duration, which indicates that closely related species that have diverged more recently tend to have more similar peak fruiting times, with a weaker but still detectable similarity in fruiting duration. This pattern likely reflects niche conservatism, whereby closely related species retain similar reproductive strategies, exposing them to shared selection pressures from frugivore availability and interspecific competition that maintain conserved peak fruiting times across evolutionary timescales (Wright and Calderon 1995; Davies et al. 2013). While our sample of 50 tree species indicates phylogenetic influences on fruiting phenology in this tree community, the strength and direction of phylogenetic control on fruiting phenology may vary across landscapes, clades, and biogeographic contexts (Davies et al. 2013). Additionally, the inclusion of functional traits that are more appropriate to phenology such as seed coat thickness, embryo size, seed viability and germination time may reveal other connections between traits and phenology. With these limitations in mind, our results nevertheless reveal a pronounced divergence in phenological control, with leaf flushing and flowering more linked to contemporaneous local climatic cues, while fruiting is less influenced by these cues and more by intrinsic, evolutionarily conserved timing mechanisms (Visser et al. 2010; Davies et al. 2013; Helm et al. 2013; Shahzad et al. 2024).

Climate projections for the southern Western Ghats indicate increasingly seasonal and unpredictable precipitation patterns alongside drier premonsoon periods, reduced annual rainfall, and warmer temperatures (Varikoden et al. 2019; Katzenberger et al. 2021). The stronger climatic sensitivity of leaf flushing and flowering across species suggests that these phenophases will show increased variability and potential desynchronization, tracking stochastic dry periods and erratic rainfall events that could trigger out-of-season leaf flushing and flowering as climate becomes more unpredictable (Chmielewski and Rötzer 2001; Hidalgo-Galvez et al. 2018; Buonaiuto and Wolkovich 2021). In contrast, the strong phylogenetic structuring observed in timing of fruiting phenology could result in a reduced capacity to track rapid precipitation and temperature change (Davis et al. 2010; Visser et al. 2010; Buckley and Kingsolver 2012; Shahzad et al. 2024). Families such as Lauraceae, Sapotaceae, Myrtaceae, Myristicaceae, and Euphorbiaceae which exhibit phylogenetically conserved fruiting during the pre-monsoon period (April – May), may be particularly vulnerable as this critical temporal window is projected to face progressive moisture deficit (Varikoden et al. 2019). With limited evolutionary flexibility to shift fruiting times due to phylogenetic constraints, these families face the risk of temporal mismatches with mutualistic animals, reduced fruit set, failed seed development, and compromised seedling germination and regeneration (Rafferty and Ives 2013; Mokany et al. 2014; Rafferty et al. 2015; Renner and Zohner 2018; Visser and Gienapp 2019; Sandor et al. 2021; Sullivan et al. 2024).

The fate of tree communities in the southern Western Ghats will therefore hinge less on shifts in climate alone but rather on the resilience of species with different reproductive phenology patterns to remain coupled to dispersal and establishment. Future research needs to focus on potential fruiting or regeneration failure of specific families or genera and investigate possible climate anomalies that trigger these changes. Identification and conservation prioritization of the identified families and lineages vulnerable to climate – fruiting mismatch will be vital for anticipating shifts in regeneration dynamics and maintaining long-term forest resilience.

## Supporting information

Online Resource 1

## Acknowledgements

We are grateful to the Tamil Nadu Forest Department for long-term research permits (WL5A/035933/2016, WL5A/631/2022, Extension Permit 4/2026) and to the Field Directors, Deputy Directors, Range Officers, and staff of the Anamalai Tiger Reserve for extending their support and cooperation. We thank Rohini Nilekani Philanthropies, Rainmatter Foundation, Arvind Datar, AMM Murugappa Chettiar Research Centre, and Cholamandalam Investment and Finance Corporation Ltd for funding support. Our colleagues T. Vanidas, Kshama Bhat, Gopana Nanda Kumar, M. Ananda Kumar, and Ganesh Raghunathan provided much help with fieldwork and at the research station. We are grateful to the organisers and many speakers at the Pheno2025 conference, Brazil, for invigorating discussions.

## Author contributions

A.P.M. contributed to data collection, performed data analysis, and wrote the first draft. D.M. and T.R.S.R. contributed to study conception, funding, manuscript writing, and editing. T.R.S.R. also contributed to data analysis. D.M. contributed to project administration and resources. S.K. contributed to manuscript editing and field data collection. A.M., K.B.S., G.M., T.S., T.V., and R.R. contributed to data collection and field work.

## Data availability

Data for the study are available in Madhavan et al. (2026). [The dataset will be made publicly available along with publication.]

## Notes

**Conflict of interest** The authors declare no conflict of interest or competing interests

### Competing Interest Statement

The authors have declared no competing interest.

