## Supplementary figures and images for "Phenology of 50 tree species across 9 years in a South Asian tropical rainforest indicates complex influence of climate, traits, and phylogeny"

### Online Resource 1

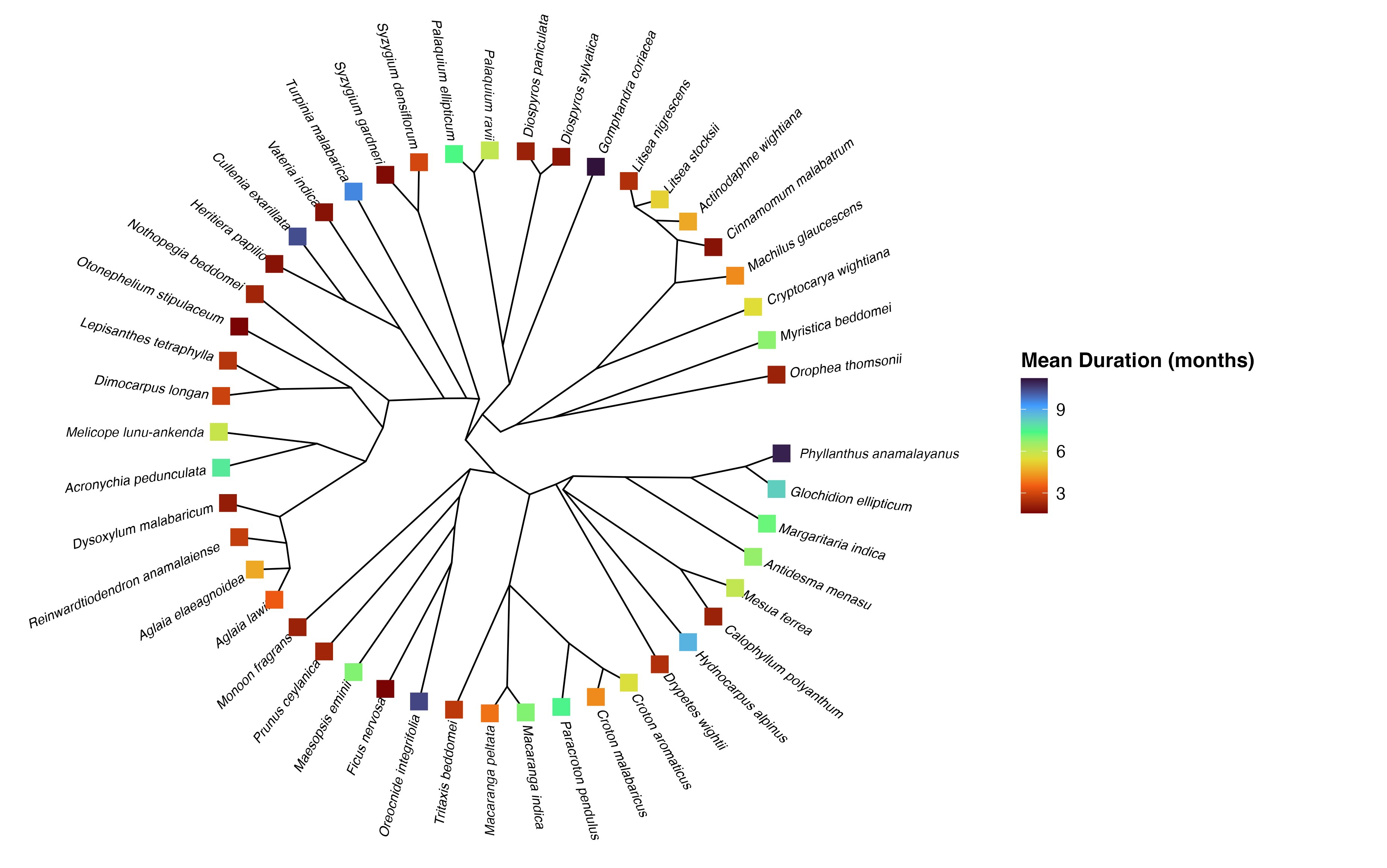
